# Chronic corticosterone enhancement aggravates alpha-synuclein brain spreading pathology and substantia nigra neurodegeneration in mice

**DOI:** 10.1101/469999

**Authors:** Johannes Burtscher, Jean-Christophe Copin, João Rodrigues, Senthil K. Thangaraj, Anass Chiki, Marie-Isabelle Guillot de Suduiraut, Carmen Sandi, Hilal A. Lashuel

**Affiliations:** Laboratory of Molecular and Chemical Biology of Neurodegeneration, Brain Mind Institute, EPFL, Switzerland; Laboratory of behavioral genetics, Brain Mind Institute, EPFL, Switzerland

**Keywords:** synuclein, Parkinson’s disease, neurodegeneration, chronic stress, amygdala, phosphorylation, behavioral, pathology spreading

## Abstract

Chronic stress and associated heightened glucocorticoid levels are risk factors for depression, a common non-motor symptom in Parkinson’s disease (PD). However, how heightened glucocorticoids influence PD neuropathology [alpha-synuclein (α-Syn) containing Lewy pathology and neurodegeneration] and disease progression is unclear. To address this knowledge gap, we investigated the impact of chronic corticosterone administration on α-Syn pathology, neurodegeneration, behavior and mitochondrial function in a mouse model of α-Syn pathology spreading after intracerebral injection of α-Syn preformed fibrils (PFFs). Our results demonstrate that heightened corticosterone aggravates neurodegeneration and α-Syn pathology spreading, intriguingly to specific brain regions, such as the entorhinal cortex. Corticosterone-treatment abolished distinct physiological adaptations after PFF-injection and induced differential physiological and behavioral consequences. Taken together, our work points to elevated glucocorticoids as a risk factor for the development of the neuropathological hallmarks of PD. Strategies aimed at reducing glucocorticoid levels might slow down pathology spreading and disease progression in synucleinopathy.

## Introduction

Synucleinopathies, such as Parkinson’s disease (PD), are diseases characterized by the accumulation and aggregation of the protein alpha-synuclein (α-Syn) and neuronal loss in the affected brain regions (neocortical, limbic and nigro-striatal circuities)^1,2^. Under physiological conditions, α-Syn is believed to play roles in synaptic transmission ^3,4^, exocytosis ^5^ and mitochondrial function ^6^. In PD brains, α-Syn undergoes conformational changes that render the protein prone to aggregation. Disease-associated mutations enhance α-Syn aggregation *in-vitro* and promote the formation of Lewy body (LB) and Lewy neuritis-like pathology in neuronal and animal models of PD ^7^. Studies using genetic manipulations of α-Syn (either knockout or overexpressing different forms of α-Syn) suggest that α-Syn misfolding and aggregation, rather than loss of α-Syn function, play central roles in the pathogenesis of PD and related synucleinopathies. This notion is supported by findings that mutations ^8–10^, duplication ^11^ or triplication ^12^ of the gene coding for α-Syn, *SNCA*, are sufficient to cause α-Syn misfolding and aggregation and early onset forms of PD. Even though overexpression of WT or disease-associated mutants of α-Syn in rodents or nonhuman primates recapitulate many pathological and motor features of PD, none of these models reproduce the full spectrum of pathological and clinical features of the disease ^13^.

Besides the cardinal motor symptoms, non-motor symptoms are common in PD. Anxiety and depression disorders for example not only often precede PD, but represent common comorbidities and non-motor symptoms of PD at later stages ^14–16^. PD-associated depression and anxiety have been linked to anatomical and metabolic alterations in the limbic system, in particular the amygdala, and the dopaminergic system ^17^. The involvement of the amygdala in stress effects, mood, emotion and reward behaviors is well-established ^18,19^, and of interest also in the context of PD. Thus, the amygdala is particularly prone to the formation of α-Syn pathology (LBs in PD-patients and LB-like pathology in many animal models)^20,21^, which occurs there as early as α-Syn pathology is observed in the substantia nigra.

Recent findings showed the spreading of α-Syn-pathology from host-tissues to mesencephalic transplants grafted into PD-patien?s brains ^22,23^ and subsequent studies provided robust evidence for inter-neuronal transmission of α-Syn-pathology ^24,25^. Motivated by these observations, several groups sought to evaluate the hypothesis that α-Syn-pathology can be induced by external seeds and be propagated through the central nervous system via a prionlike mechanism ^26–28^. This hypothesis has been tested in rodents treated with different forms of recombinant α-Syn aggregates (reviewed in ^29^) or with α-Syn aggregates derived from postmortem human brains from patients with α-Syn-pathology, PD ^30^ or MSA ^31,32^. Injection of α-Syn PFFs in different brain regions, such as the striatum ^27^, the olfactory bulb ^33^ or the substantia nigra ^28^ induce pronounced α-Syn pathology spreading, often strongest to the amygdala. Despite the observations of preferential accumulation of α-Syn-pathology in the amygdala, literature linking α-Syn-pathology spreading and amygdala-related behavior or physiology is sparse.

Chronic stress and associated heightened glucocorticoid levels are known risk factors for anxiety and depression ^34,35^, and have also been suggested to be risk factors for neurodegeneration in mouse PD models ^36,37^. Therefore, we sought to investigate whether chronic elevation of corticosterone (CORT) would aggravate α-Syn pathology spreading and neurodegeneration after intrastriatal injection of α-Syn PFFs. CORT was delivered in the drinking water over a period of 11 weeks to mice, a regime known to increase depression-like behaviours ^38–40^. We then explored the possibility that chronic heightening of CORT and α-Syn pathology synergistically increase behavioral deficits related to motor and non-motor symptoms of PD. Systemic administration of CORT has been demonstrated to increase the activity of the basolateral amygdala ^18^ and depression in PD is associated with increased metabolic activity in the amygdala ^17^. Therefore, we also studied the metabolic and behavioral consequences of α-Syn pathology in the amygdala.

## Results

### Experimental design and rational

We generated PFFs from recombinant wild-type α-Syn for intrastriatal injections in mice (suppl. Fig. 1). Analyses by transmission electron microscopy (TEM) revealed the presence of fibrils of different morphologies (suppl. Fig. 1a), including straight and twisted PFFs, resembling the heterogeneity of α-Syn fibrils observed in human LB pathology ^41^. As expected, the PFFs bound the amyloid-specific dye Thioflavin T and after sonication existed predominantly as short fibrils with a median length of 80 nm (suppl. Fig. 1b and 1c). The PFF preparations contained mainly fibrils, and 10-20 % monomeric α-Syn (suppl. Fig. 1d) that is generated during sonication and is in equilibrium with the fibrils through constant recycling of the monomers at the fibril ends ^42^. This level of monomers was maintained in our preparations to enhance amyloid formation and the seeding capacity of the PFFs post injection ^43^.

Our behavioral analyses focus on a time interval of 1-2 months after intrastriatal α-Syn PFF injection. In our experiments, we consistently observe a peak of α-Syn pS129-positive aggregate levels in the amygdala and cortical regions between 1 to 3 months after intrastriatal α-Syn PFF injection (Fig. 1a). The main brain regions of interest were the striatum (site of PFF injection), the substantia nigra (neurodegeneration in which is crucial for cardinal motor symptoms in PD) and the amygdala (high pathology spreading). In the striatum, α-Syn pS129 related pathology is predominantly neuritic 1 month after injection, whereas more peri-nuclear, compact, often half-moon shaped inclusions are observed at later time points (Fig. 1b, c), without observable loss of pS129 signal. To investigate the relationship between α-Syn pathology and potential brain region related behavioral deficits in the time of highest α-Syn aggregation load in the amygdala and other brain regions, behavioral tests were conducted 1-2 months after PFF injection.

**Fig. 1:**
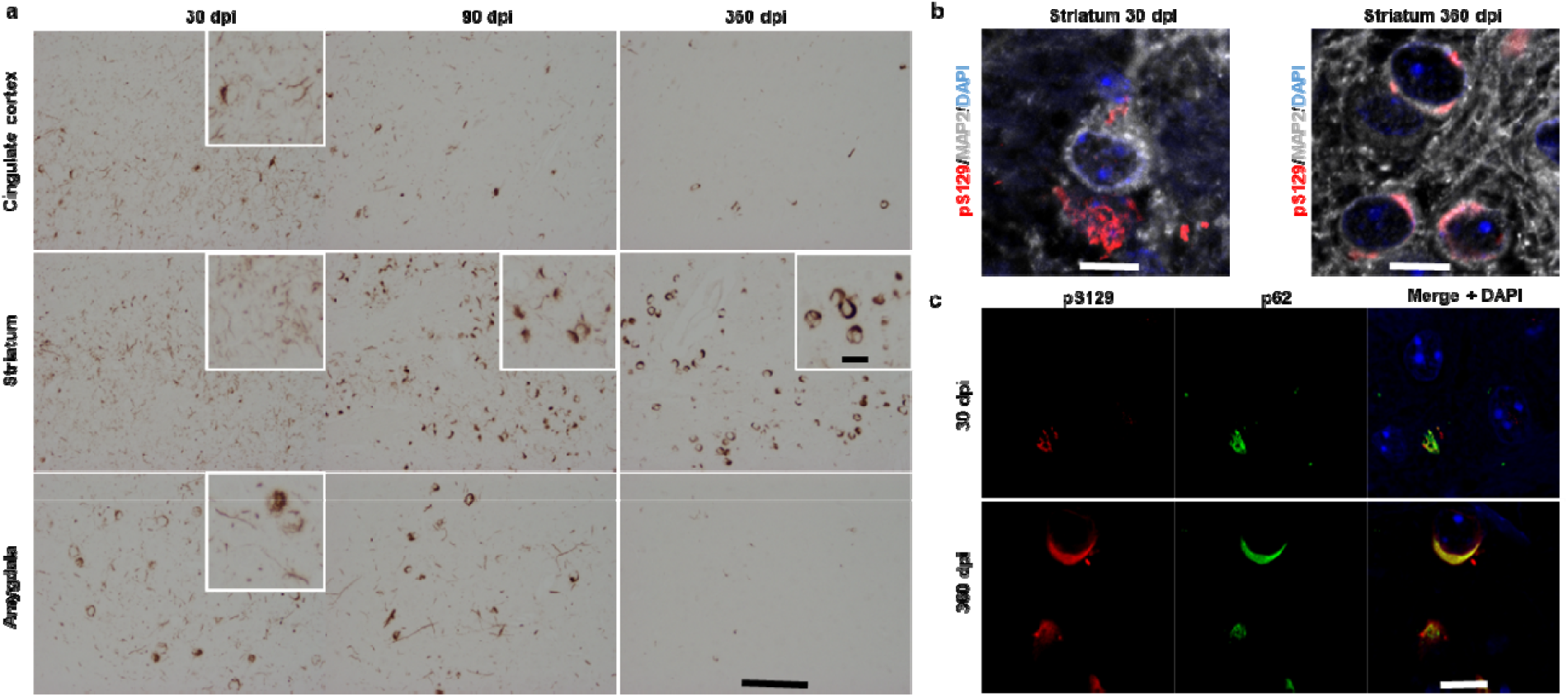
Evolution of pS129 a-Syn pathology in the cingulate cortex, striatum and amygdala over time. (a) α-Syn pS129 immunoreactivity in the cingulate cortex, striatum and amygdala (each in the hemisphere of injection) 30, 90 and 360 days post injection (dpi) after unilateral, intrastriatal injection of PFFs. (b) Striatal MAP2-positive (grey) neurons stained for α-Syn pS129 (red) and at 30 and 360 dpi and (c) co-stained for α-syn pS129 and p62. Scale bars are 100 μm (a), 20 μm (inset) and 10 μm (b and c).

Young adult, male mice were treated with CORT or vehicle, after which they were injected unilaterally with either α-Syn PFFs [hereafter referred to as PFF(C) mice] or PBS [PBS(C) mice] into the dorsal striatum by stereotactic surgery. Vehicle treated mice are referred to as PBS-mice or PFF-mice, respectively. After surgery, CORT/vehicle treatment was continued until sacrifice (Fig. 2a). Half of the animals were used to assess mitochondrial parameters one month after PFF injection, the other half were subjected to behavioral testing followed by brain processing for histological analyses.

**Fig. 2:**
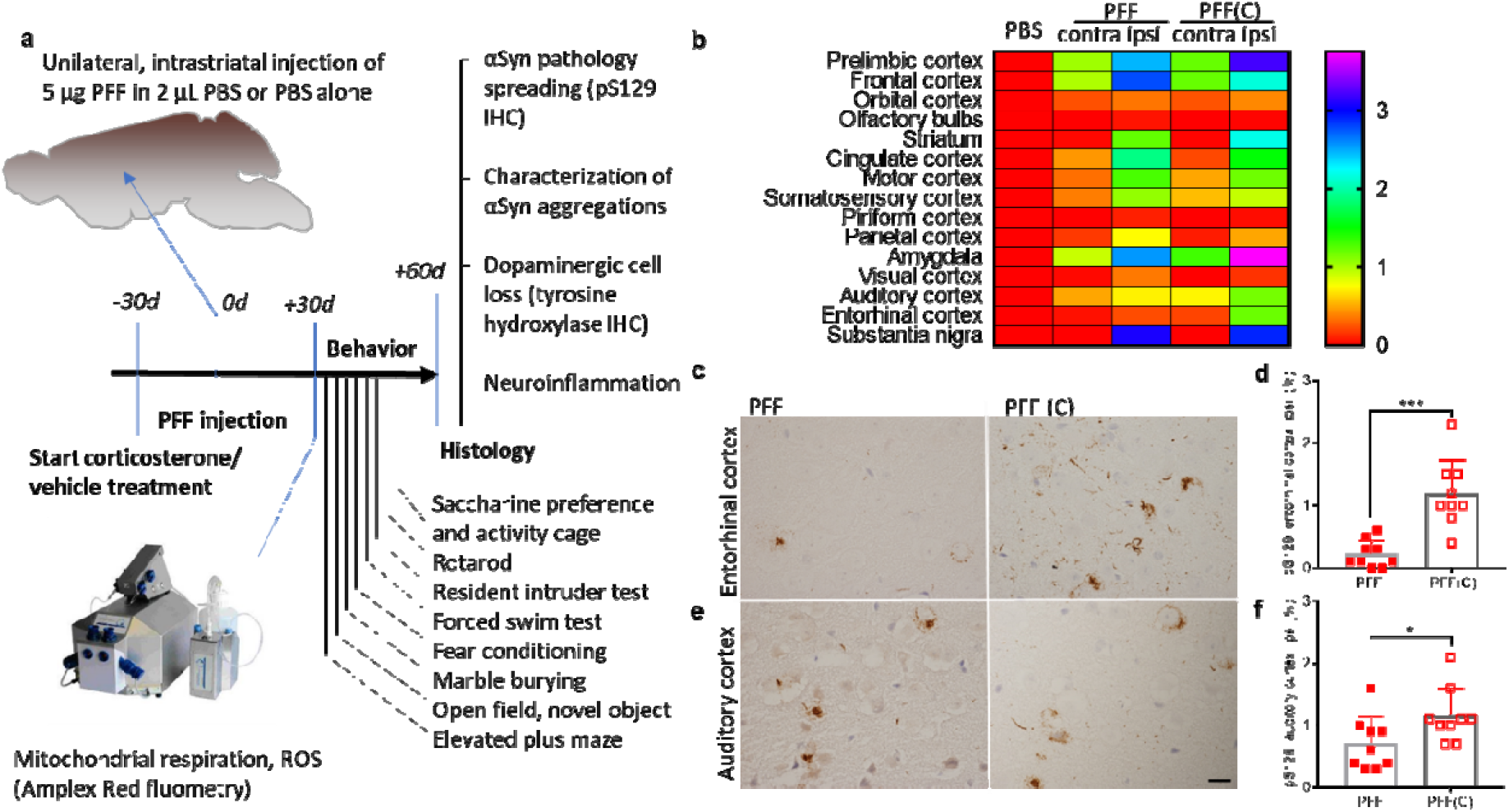
Experimental setup and α-Syn spreading pathology. (a) Experimental setup: 5 μgof PFFs in 2 μL PBS (or PBS alone) were injected unilaterally in the dorsal striatum of mice that were continuously treated with either corticosterone (35 mg/L) or vehicle (0.45% hydroxypropyl-β-cyclodextrin) in the drinking water throughout the experiment, starting 1 month before PFF/PBS-injection. 1 month after injections, some animals were sacrificed for high-resolution respirometry experiments and the other group of animals was subjected to a series of behavioral tests lasting for about another month, after which brains were processed for histology (IHC = immunohistochemistry). (b) Heat map reflecting mean α-Syn pathology spreading across the brain, ipsi(lateral) and contra(lateral) (density of α-Syn pS129 immunoreactivity in % of area). Significant differences in α-Syn pathology spreading were observed in 2 brain regions; entorhinal cortex (c,d; Mann-Whitney test, P<0.001) and auditory cortex (e,f; Mann-Whitney test, P<0.018) between corticosterone- [PFF(C)] vs. vehicle- [PFF] pretreated animals. Scale bar represents 20 μm. N=9 mice per group.

Treatment with CORT (Fig. 2a, suppl. Fig. 2a) induced depressive like phenotypes in the forced swim test (FST, suppl. Fig. 2b,c and 3) and in a saccharine preference test (suppl. Fig. 2d), indicating anhedonia-like behavior. Chronic CORT furthermore had pronounced effects on weight gain after surgery and body fat content normalized to body weight (suppl. Fig. 2e,f,g), a well-known effect of elevated CORT-levels^39,44^ as well as on drinking and feeding behavior (suppl. Fig. 2h,l; note that CORT treatment was discontinued during activity measurements in the home cage, in which feeding and drinking behavior were dramatically changed). 2-way ANOVAs revealed significant effects of CORT treatment in the absence of PFF influences and interaction effects in all these parameters.

### CORT-treatment aggravates α-Syn pathology spreading and dopaminergic cell loss in α-Syn PFF-injected mice

We then investigated, whether the pathological hallmarks of PD – α-Syn/Lewy-pathology and dopaminergic cell loss – are influenced by heightened corticosterone upon injection of PFFs. α-Syn pathology spreading was assessed histologically by densitometry of α-Syn pS129 immunoreactivity 60 days after PFF ? PBS injection (Fig. 2b). No α-Syn pS129 immunoreactivity was detected in PBS-injected controls. In PFF-injected animals, the highest α-Syn pS129 densities were observed in the hemisphere of injection in the amygdala, prelimbic cortex and substantia nigra for both, CORT and vehicle treated groups (Fig. 2b). α-Syn pS129 signal in the entorhinal cortex was almost absent in the PFF group and significantly higher in the PFF(C) group. In the auditory cortex, α-Syn pS129 density was significantly higher in PFF(C) mice (Fig. 2c). No significant differences in spreading were observed in other brain regions (suppl. Fig. 4). 60 days after PFF-injection, no dopaminergic cell loss in the substantia nigra was observed in PFF mice. On the other hand, CORT treatment resulted in decreased density of tyrosine-hydroxylase (TH) immunoreactivity and reduced ipsilateral (hemisphere of injection) to contralateral TH-positive cell numbers in the hemisphere of PFF-injection as compared to the contralateral substantia nigra (Fig. 3a-d); no such effects were observed in PBS-injected groups (suppl. Fig. 5a-c). The density for α-Syn pS129-positive aggregates (Fig. 3e,f) and colocalization of pS129 with TH (suppl. Fig. 5d) were similar between PFF and PFF(C) animals. α-Syn pS129-positive aggregates colocalized with the macro autophagy marker p62 and ubiquitin in in the substantia nigra of PFF and PFF(C) mice (suppl. Fig. 6), and in other brain regions (suppl. Fig. 7). The aggregates were resistant to proteinase K treatment and were detected by antibodies for pS129 and for the N-terminal part of α-Syn (1-20) (Fig. 3g). Motor coordination in the rotarod test was significantly reduced in PFF(C) mice, but was mainly due to CORT treatment, and not due to PFF or interaction (suppl. Fig. 5f). Total distance travelled during 3 days in the activity cage (AC) was similar across groups (suppl. Fig. 5e). Altogether, these results suggest that chronic CORT-treatment aggravated substantia nigra neurodegeneration after PFF injection without affecting pS129 immunoreactivity quantitatively (Fig. 3f) or qualitatively (Fig. 3g and suppl. Fig. 6), general motor behavior in the AC or motor coordination in this time interval.

**Fig. 3.**
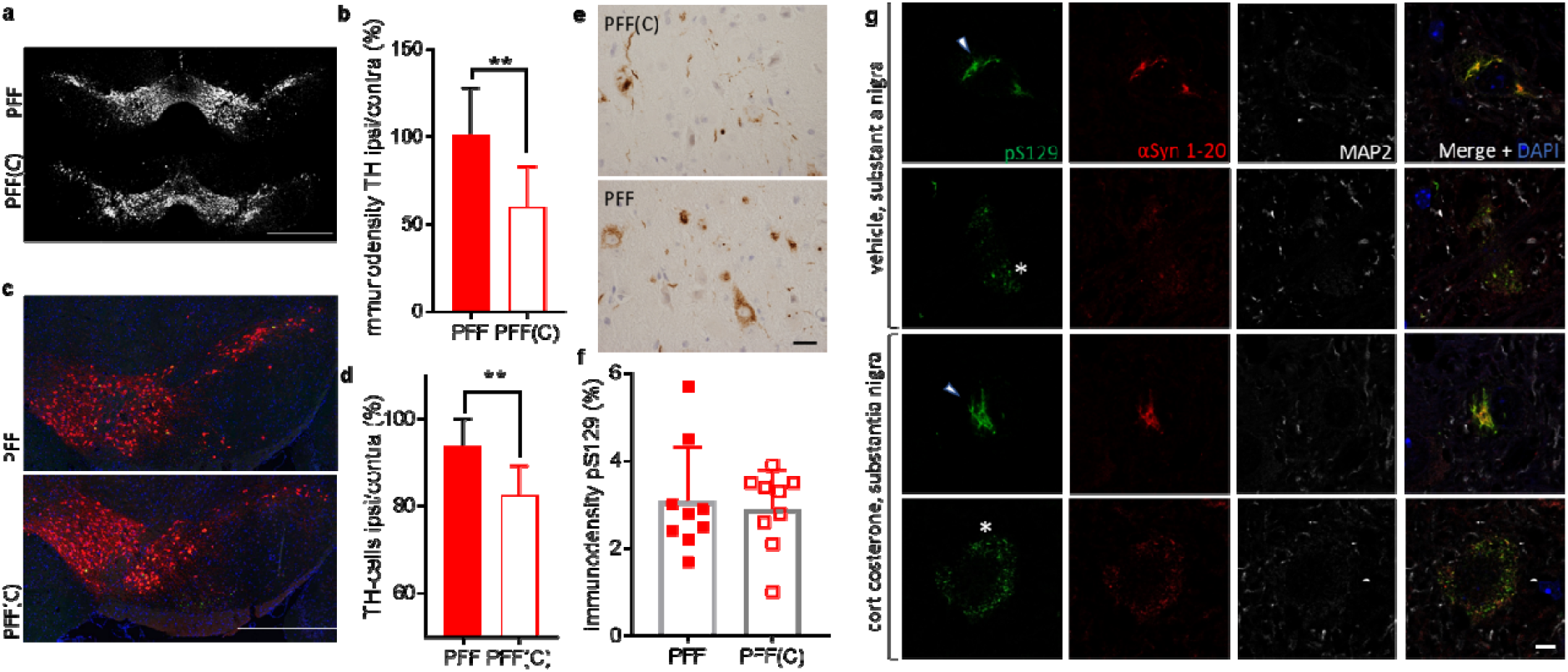
α-Syn preformed fibril (PFF) injections aggravated neuropathology in the substantia nigra after chronic corticosterone (C). (a) Depictions of tyrosine hydroxylase (TH)-labelled representative midbrains, including the substantia nigra and (b) quantification of densities of TH immunoreactivity (p=0.003, t-test). (c) Micrographs of ipsilateral (i.e. hemisphere of injection) midbrains including the substantia nigra, pars compacta and pars reticulata (TH in red, pS129 in green and DAPI in blue) and (d) cell counts of ipsilateral substantia nigra tyrosine hydroxylase positive neurons normalized to contralateral cell count (p=0.002, t-test). Values reflect mean cell counts of 6 sections per brain each, (e) α-Syn pathology spreading (pS129 density) in the substantia nigra and quantification (f). (g) characterization of aggregates in the substantia nigra using immunostaining for pS129, α-Syn 1-20 and microtubule associated protein 2 (MAP2) after proteinase K treatment. Note the two clearly distinguishable aggregation forms: fibrillar (arrowheads) and granular (asterisks). Scale bars represents 500 μm (a,c), 20 μm (e) and 5 μm (g). N=9 per group for all quantifications.

### CORT-treatment reverses anxiety-effects of α-Syn PFF injection

Elevated CORT has been shown to affect mood and emotional behavior ^38–40^, and Chronic CORT also affects depressive-like behavior, independently of PFF injection (Fig. 2). Given our observation of pronounced α-Syn pathology in several brain regions implicated in mood and emotional behavior, such as the amygdala (Fig. 1, 2), we also investigated anxiety-like behaviors. We assessed basal spontaneous anxiety and exploratory reactivity to novelty in the elevated plus maze (EPM) and open field test (OF). The OF was followed by a novel object test, assessing anxiety-linked novelty seeking. No differences in all these anxiety parameters were observed between PBS-injected groups (suppl. Fig. 8a-i). However, we observed a moderate hypo-anxious phenotype for PFF mice in the EPM, which was reversed by CORT-treatment (Fig. 4a-f). PFF(C) animals also moved less in the EPM (Fig. 4d). In the OF, a similar effect was observed (Fig. 4g,h): PFF-injection induced hypo-anxious phenotypes (more time spent in the center and less time spent at the walls of the arena), which was reversed by chronic CORT-treatment, resulting in statistically different behavior in the OF between the PFF-injected groups. The mice in all groups moved at similar speed in this test (Fig. 4i). Neither CORT-treatment, nor PFF-injection significantly impacted on the interest in a novel object (suppl. Fig. 8j-m). A marble burying test, used to assess neophobia and anxiety, revealed no differences across groups (suppl. Fig. 9a,b). Taken together, PFF-injection resulted in a moderately hypo-anxious phenotype, which was reversed by CORT-treatment, in the EPM even trending towards hyper-anxious phenotypes.

**Fig. 4.**
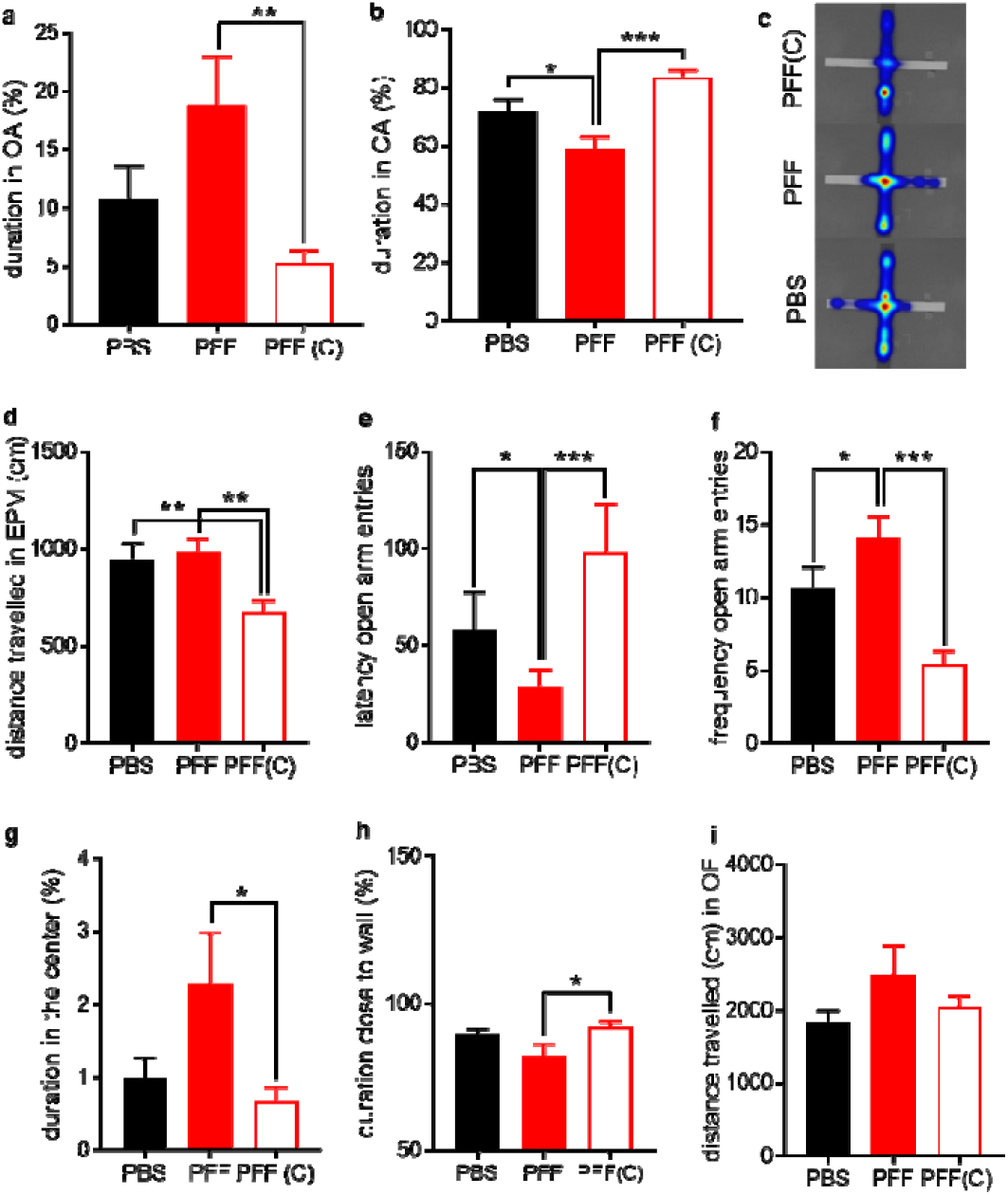
Corticosterone (C) treatment reversed preformed fibril (PFF) induced anxiety-like behaviors. A hypo-anxious phenotype in the elevated plus maze (EPM) of PFF-injected mice was reversed by C-treatment, in the open arms (OA, a) and the closed arms (CA, b): lway ANOVAs, F (2, 51) = 5.384, P=0.008 and F (2, 51) = 12.17, P open field test (OF), mice injected with PFFs spent more time in the center, if not treated with C (F (2, 24) = 3.604, P=0.043). Mice treated with C and PFFs accordingly spent more time close to the walls (h), indicating higher anxiety-like behavior with C treatment: lway ANOVA, F (2, 24) = 3.464, P=0.048. No differences between groups were observed for distance travelled in the OF. N=17-18 per group for a-f, N=9 per group for g-i.

### α-Syn pathology in the amygdala does not alter mitochondrial respirational capacities, fear behavior or aggression

We observed the highest levels of α-Syn pathology of all brain regions in the ipsilateral amygdala (hemisphere of injection) of PFF(C) animals (Fig. 2b). α-Syn pS129 levels, assessed by immunostaining (Fig. 5a,b) were not significantly different in the amygdala of PFF and PFF(C) mice. We hypothesized that the aggregation load in the amygdala should negatively impact on its physiology. Due to the involvement of the amygdala in fear-related and aggressive behavior ^45,46^, we assessed potential impairment of these behaviors applying a fear conditioning protocol and a resident intruder aggression test. As monomeric and aggregated α-Syn species have been shown to affect mitochondrial function, we hypothesized that α-Syn pathology might disturb mitochondrial functions in the amygdala. More specifically, the mitochondrial import machinery ^47^, mitochondrial membrane potential, cytochrome c release, ROS and morphology ^48,49^ as well as mitochondrial permeability transition pore and electron transfer system regulation ^6,50^ have all been implicated to be affected by α-Syn. We observed, however, no adverse effect of amygdala α-Syn pathology on mitochondrial functions *in vivo*. Mitochondrial respiration (Fig. 5c, suppl. Fig. 10), mitochondrial ROS-production (Fig. 5d, suppl. Fig. 10), flux control ratios (suppl. Fig. 11e) and oxidative phosphorylation coupling efficiency (respiratory control ratio, RCR) (suppl. Fig. 12b) did not differ across groups (controls and contralateral data in suppl. Fig. 10). Notably, at the site of PFF-injection (striatum), we observed a significantly higher reliance of mitochondrial respiration on the succinate pathway of PFF mice as compared to PFF(C) mice (suppl. Fig. 13e,f). PFF and PFF(C) mice also did not behave differently compared to PBS-injected controls in fear conditioning (Fig. 5f,g and suppl. Fig. 14) and resident intruder tests (suppl. Fig. 15).Thus, α-Syn pathology does not appear to significantly impair amygdala mitochondrial physiology or amygdala-related behaviors.

**Fig. 5.**
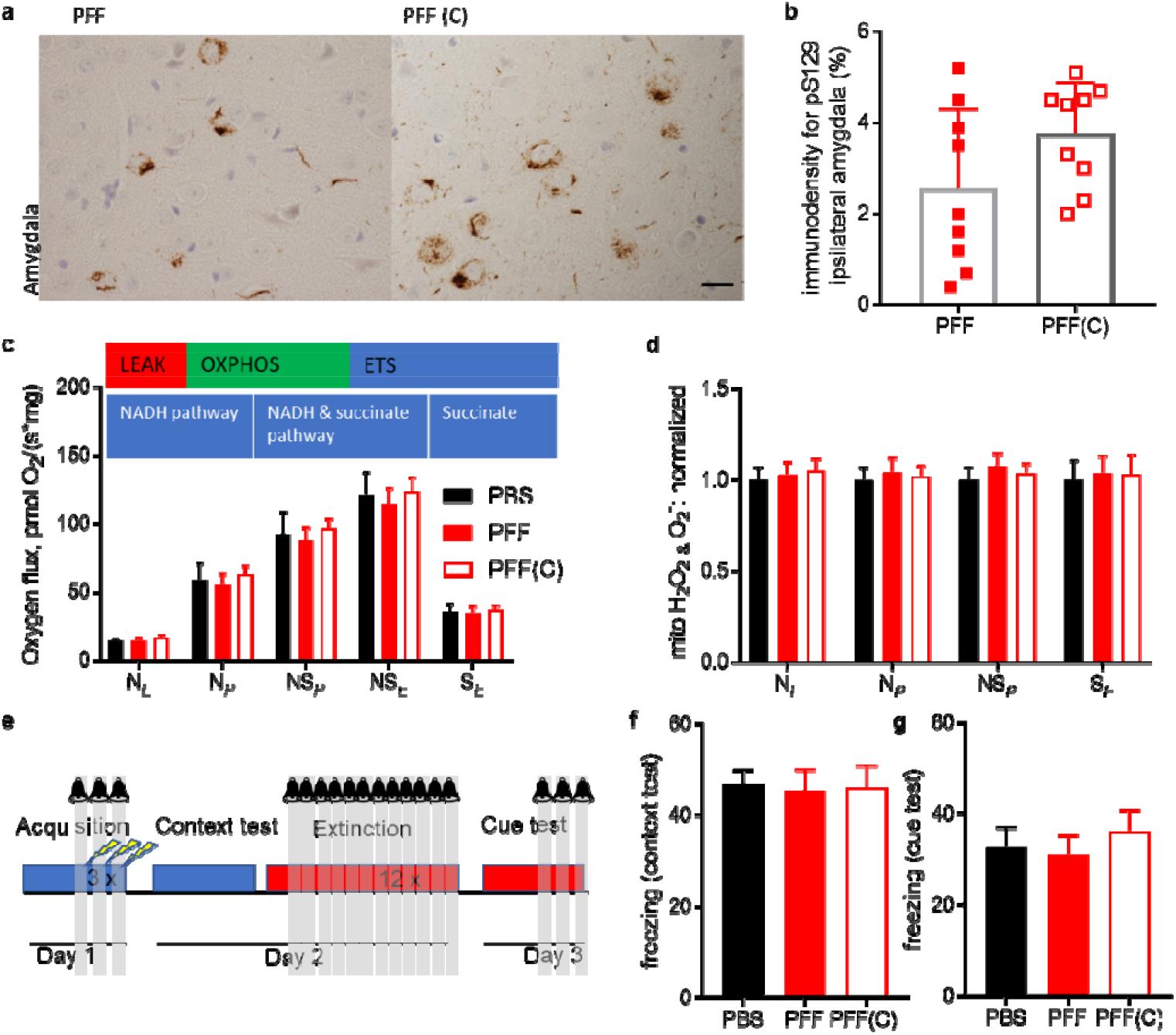
α-Syn pathology in the amygdala after PFF-injection does not affect mitochondrial function and amygdala-related behaviors. (a) Similar α-Syn pS129 immunoreactivities in the basolateral amygdala (scale bar represents 20 μm) are quantified in (b); P=0.14, Mann-Whitney. Mitochondrial respiration was assessed in LEAK and OXPHOS states driven by the NADH pathway (N¿ and N_P_), in the OXPHOS state driven by NADH and succinate pathway combined (NS_P_), the electron transport system (ETS) state driven by NADH and succinate pathway combined (NS_f_) or only by the succinate pathway (S_f_). No differences in respiration were observed in any state across the groups (c). Amplex red fluorometry performed in parallel to respiration revealed no differences in mitochondrial hydrogen peroxide or superoxide production in any state (d): ROS-levels are normalized to PBS-injected controls (no corticosterone (C) treatment). Neither α-Syn pathology, nor additional CORT treatment affected fear related behaviors (e-g). Scheme of fear conditioning experiments (e), freezing behaviors in the context (f) and in the cue (g) test. N=8-9 per group.

### Discriminant analysis reveals differential effects of α-Syn PFFs in heightened CORT conditions

To investigate in more detail the relations of α-Syn, CORT-treatment and behavioral components, partial least squares (PLS) discriminant analysis was performed (Fig. 6). The 6 variables with highest variable importance in projection (VIP) for each component used in a 2-component model (Fig. 6a) were identified (body fat content, weight gain, behavior in the FST, as well as drinking, feeding and hedonic behavior in the AC), yielding good predictive capacity for the CORT condition (Fig. 6b). The heatmap in 6c depicts the relatively faithful prediction of VIP variables, and scatterplots in 6d reveal good separation of the PFF(C) mice from other treatment groups.

**Fig. 6:**
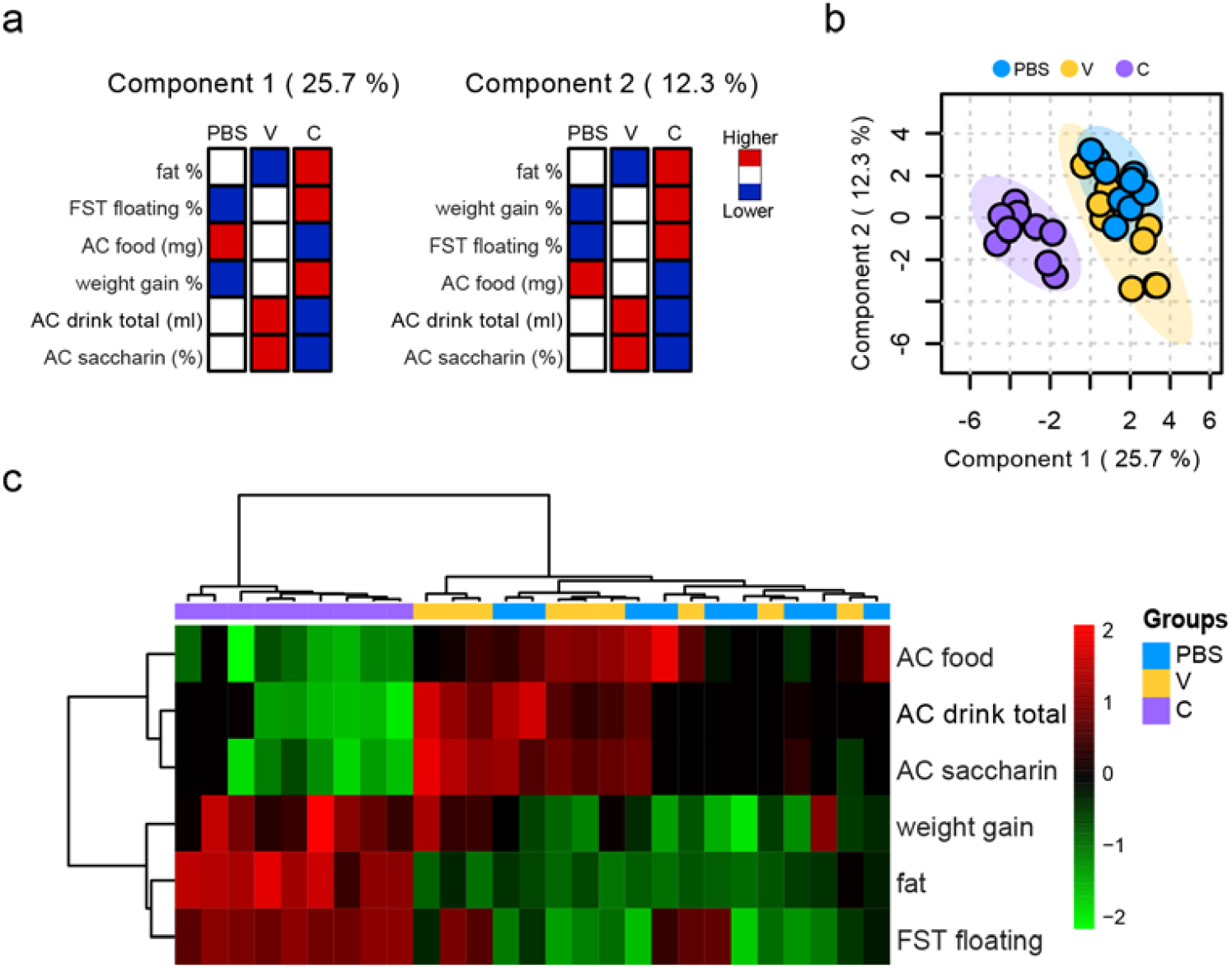
PLS Discriminant analysis results. Model accuracy was used to decide for the number of components. Two components were chosen, resulting in a model accuracy of approximately 76 %. Empirical p-value for using 2 components, estimated with permutation statistics (N permutation = 2000, p=0.004). (a) the 6 variables with highest variable importance in projection (VIP) for each component used in the model and their respective contribution for each experimental group: “fat” is the percentage of fat per body mass, “FST floating” is the percentage of time in the FST (forced swim test) spent in immobility, “AC drink total” are consumption (water and saccharine solution combined, in ml) in the AC (activity cage), “AC saccharin” describes the percentage of saccharin solution consumed per total consumption of water and saccharine solution combined, “AC food” is the total food consumed during the AC. (b) Scatterplot depicting values of individual mice for the 2 components, (c) heat map of the previous variables for each subject (each column summarized data of 1 animal). Corticosterone (C) and vehicle (V) pretreated mice injected with PFFs and the PBS-injected control group are compared.

### Correlation of α-Syn pathology with behavioral phenotypes

Next, we sought to investigate, whether brain-region specific α-Syn pathology correlated with the performance in behavioral tests. We observed several significant correlations (Fig. 7). Due to the absence of α-Syn pS129 immunoreactivity in PBS-injected controls, only PFF-injected animals were subjected to correlation analyses. CORT and vehicle pretreated cohorts exhibited differential correlation patterns. For example, substantia nigra α-Syn pS129 density correlated to several behavioral parameters in the CORT, but not the vehicle group (suppl. Fig. 16).

**Fig. 7.**
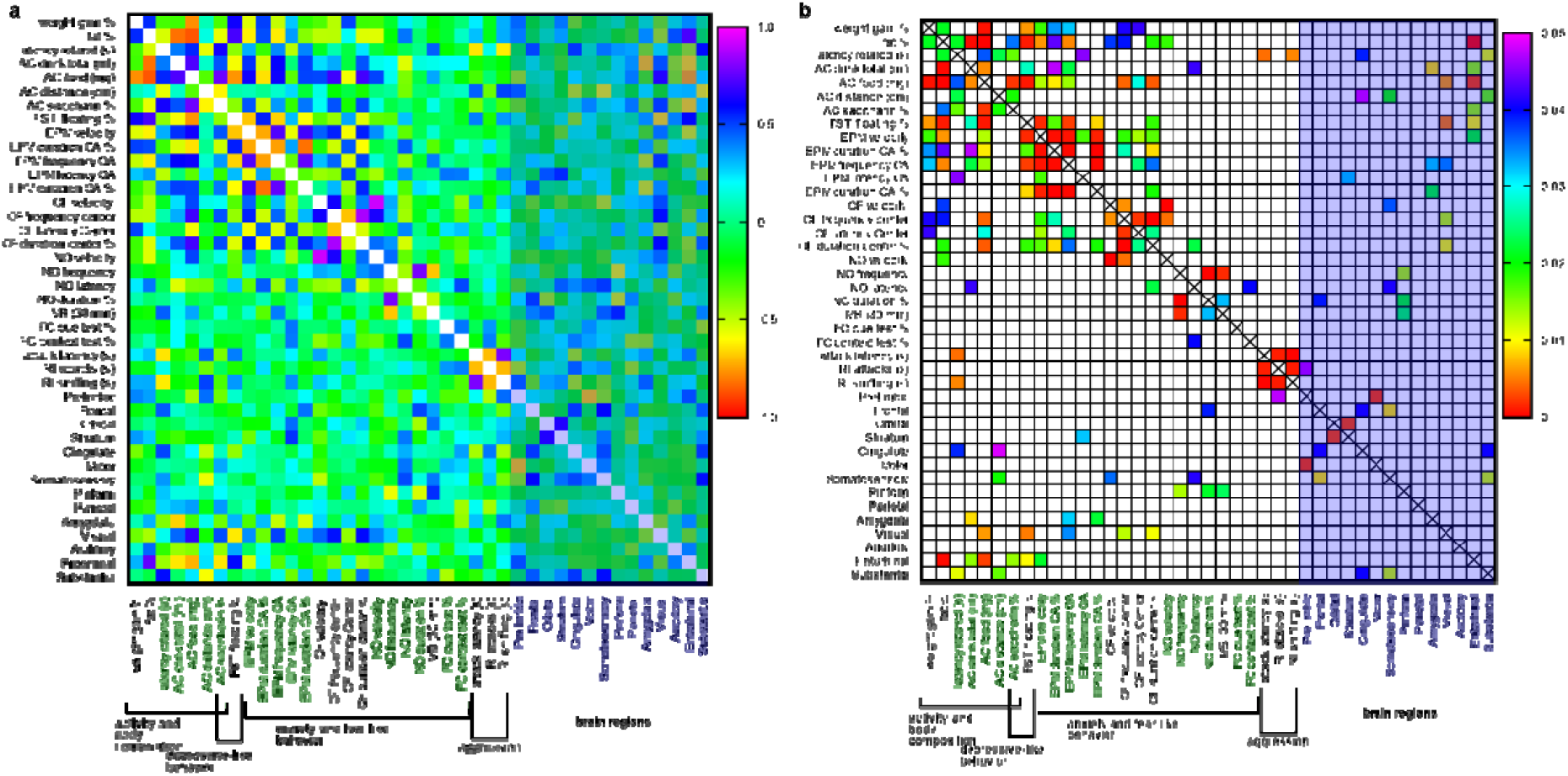
Correlations between behavioral measures and α-Syn pS129 immunoreactivity in specific brain regions of animals injected with α-Syn preformed fibrils (corticosterone and vehicle pretreated cohorts pooled). Pearson’s correlation coefficients are represented in (a), while according P-values (Á0.05) are indicated in (b). AC = activity cage, FST = forced swim test, EPM = elevated plus maze, OF = open field test, NO = novel object test, MB = marble burying test, FC = fear conditioning, Ctx = cortex

We applied partial least squares (PLS) regression analysis, followed by multivariate ANOVA to further elucidate the dependent behavioral variables predicting α-Syn pathology in different brain regions (Fig. 8). Mean values of α-Syn pS129 immunoreactivity (% of area) for selected brain regions (PFF and PFF(C) mice pooled) were used to select additional brain regions with high α-Syn pathology (8a), beside regions of main *a-priori* interest (striatum, substantia nigra, amygdala) and the entorhinal cortex that emerged of being most affected by CORT-treatment after PFF-injection. Components for all PFF-injected mice (PFF and PFF(C) mice pooled) were calculated and used to extract differences between PFF and PFF(C) mice (8b). The resulting models explained 15-25 % of variability for several selected brain regions (Fig. 8c, upper panel). Scores of behavioral and physiological outcomes on the components are depicted in Fig. 8c (lower panel). Interpretation example: striatum component 1 (Fig. 8c) explains that α-Syn pathology of the striatum was associated with lower values of consumed food in the AC, saccharine preference in the AC, velocity in the EPM, and in the OF. While PFF(C) mice had positive scores on striatum component 1, PFF mice scored negatively (Fig. 8b). Therefore, the CORT intervention impacted negatively on these striatum-related behaviors.

**Fig. 8:**
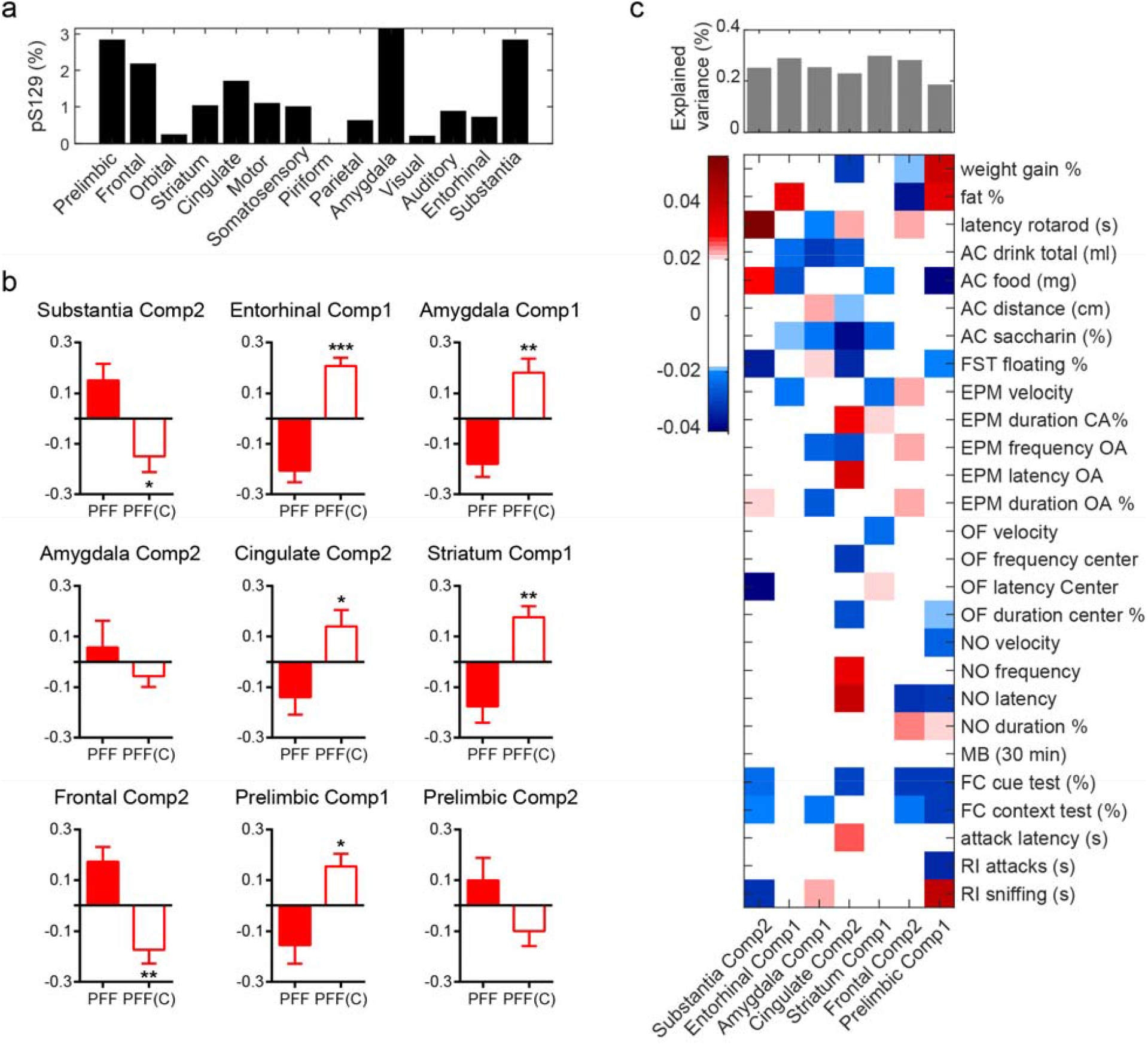
PLS Regression components with significant group differences on PFF-injected animals. a) Mean values of α-Syn pS129 immunoreactivity (% of area) for selected brain regions; b) Group differences [PFF vs. PFF(C) groups] for PLS component (Comp) values with percentage of variance explained larger than 15%. ANOVA tests following a significant MANOVA (P=0.003) with P-values corrected for multiple comparisons with the Holm–Bonferroni method; c) percentage of variance explained (top bar plot) and weights (bottom matrix) of the PLS regression components for each brain region where there was a significant effect of Group. AC = activity cage, FST = forced swim test, EPM = elevated plus maze, OF = open field test, NO = novel object test, MB = marble burying test, FC = fear conditioning, Rl = resident intruder test Significance levels: * *p* < 0.05; ** *p* < 0.01; *** *p* < 0.001

## Discussion

Despite profound impact on quality of patient life ^51^, non-motor symptoms, including affective disorders, remain less well characterized compared to motor symptoms and their impact on PD pathology and disease progression remains unknown. Several previous studies in rodent models of PD fortified the link with affective disorders ^52 53^. Chronic mild stress induced depression was also shown to worsen neurochemical and behavioral outcomes in the MPTP (1-Methyl-4-phenyl-1,2,3,6-tetrahydropyridin) model of PD ^54^. Chronic stress and stress-associated heightened glucocorticoid levels are known risk factors for these affective disorders ^34,35^, and potential risk factors for neurodegeneration, as has been demonstrated in mouse PD models ^36,37^. Despite the associations between glucocorticoid levels, affective disorders and PD, the relationship of models of (depression-inducing) heightened glucocorticoid and α-Syn pathology spreading has not been assessed previously. On the bases of these observations, we hypothesized that heightened CORT levels could influence α-Syn pathology and neurodegeneration in PD and sought to test this hypothesis in the well-established mouse model of α-Syn pathology spreading induced by intrastriatal injection of α-Syn PFFs.

We report aggravated α-Syn pathology spreading and neurodegeneration at chronic CORT administration and after PFF injection. These effects coincided with differential behavioural effects as compared to PFF-injected mice without CORT treatment.

### CORT-treatment aggravates α-Syn pathology spreading and dopaminergic cell loss in α-Syn PFF-injected mice

Unilateral injection of α-Syn PFFs into the striatum caused pronounced pathology spreading to various brain regions two months later. α-Syn pathology spreading was aggravated in distinct brain regions after CORT pretreatment, most notably in the entorhinal cortex. Interestingly, the entorhinal cortex is severely affected by Lewy pathology in many PD ^55,56^ and dementia with LB ^57^ patients. We observed that the entorhinal cortex α-Syn pathology was significantly correlated with several behavioral measures assessed, including depressive-like behavior in the FST, motor behavior in the EPM and saccharine preference. Our results point to the entorhinal cortex as being a particularly vulnerable brain region for α-Syn pathology in conditions of glucocorticoid dysbalance. α-Syn pathology in this region after PFF-injection of vehicle-pretreated animals was sparse or absent, but considerable pS129 pathology was observed in all PFF(C) mice. Due to the entorhinal cortex’ prominent role in cognition and potential role of entorhinal Lewy pathology in cognitive deficits in PD ^55,58^, studies on PD patients assessing the effect of chronic stress and depression on cognitive performance and α-Syn pathology in the entorhinal cortex will be of interest.

No neuronal loss in the substantia nigra was observed in PFF mice, which is in line with previous reports, in which neurodegeneration is detected only ∼ 180 days after injection ^27^. PFF(C) mice presented with reduced tyrosine hydroxylase (TH) immunoreactivity and a decreased ratio of TH-positive neurons in the substantia nigra (pars compacta & reticulata) of the hemisphere of injection as compared to the contralateral hemisphere. α-Syn pS129 pathology in the substantia nigra was not different between CORT or vehicle treated mice. These findings demonstrate that heightened CORT-levels aggravated α-Syn pathology spreading and nigral neurodegeneration following intrastriatal PFF injection.

### CORT-treatment reverses anxiety-effects of α-Syn PFF injection

Chronic CORT treatment induced depressive-like phenotypes and pronounced physiological changes, independent of PFF injection. CORT treatment surprisingly reversed hypo-anxious-like behavior induced by α-Syn PFF injection in the EPM and OF. This finding seems particularly relevant as hypo-anxiety is a common observation in models of early stages of PD ^59,60^, and might reflect changes in dopamine signalling ^61^. We speculate that chronic CORT treatment impaired dynamic adaptations to α-Syn pathology, thereby preventing potential transient bursts in dopamine signaling, that might occur due to α-Syn’s suspected function as negative regulator of dopamine release ^4,62^. In line with this assumption is a recent report on enhanced presynaptic activity of neurons in presence of α-Syn inclusions ^63^. Such dynamic adaptations might comprise shifts in the contributions of mitochondrial respiration pathways, as observed in the striatum. PFF animals exhibited increased relative contribution of the succinate pathway to overall mitochondrial respiration as compared to PFF(C) mice. Impairment of succinate dehydrogenase activity in the striatum has been linked to excitotoxicity ^64^ and protocols of succinate dehydrogenase inhibition (“chemical preconditioning”) have been shown to induce neuroprotection ^65^, suggesting its involvement in neuroprotective adaptations following challenges. Therefore, our findings support the view that α-Syn pathology at early time points (1-2 months) after injection of PFFs is associated with adaptive changes in the striatum. Such adaptations could include a shift of mitochondrial respiration towards (potentially protective) stronger reliance on succinate- / mitochondrial Complex II-linked respiration (S_f_), which is blocked by chronic CORT. Interestingly, the effect size of *S_E_* FCR between PFF(C) and PFF groups is comparable to the differences observed in anxiety-like behaviors, behaviors with important striatal participation ^66^.

In summary, we report that alterations in anxiety- and depression-like behaviour due to α-Syn pathology were not exacerbated by our heightened CORT treatment. Whereas depressive-like phenotypes were solely attributable, in a non-additive manner, to CORT treatment, CORT reversed PFF-induced changes in anxiety-like behaviour in some tests (EPM, OF).

### α-Syn pathology in the amygdala does not alter mitochondrial respirational capacities, fear behavior or aggression

A high level of α-Syn pathology in the amygdala has been reported in patients suffering from PD and other neurodegenerative diseases^21^, as well as in several α-Syn pathology spreading models ^27,28,30,33^. The amygdala is importantly involved in emotional behavior and depression in general^19^ and in PD in particular^51^. Therefore, we sought to determine, whether chronic CORT treatment would aggravate associated symptoms in a widely used α-Syn pathology spreading mouse model, in particular related to the amygdala.

We observed a similar strong pathology spreading to the amygdala following PFF injection, irrespective of CORT treatment. Despite the severe α-Syn pathology observed in PFF-treated animals, several amygdala-related behaviors (e.g., fear-related behaviors, aggression) were unaffected. Furthermore, PFF treatment did not affect several mitochondrial parameters measured in the amygdala. Taken together, these results suggest that α-Syn aggregations by themselves are not immediately toxic. This finding is in line with recent observations that hippocampus-dependent behavior is not altered by the induction of severe hippocampal α-Syn pathology either^67^. Alternatively, it is possible that, at the time points following treatments at which the current study took place, the relevant neuronal circuits are resilient or plastic enough to prevent general physiological deterioration or behavioral alterations. Over time only some particularly vulnerable neuronal populations – such as TH-neurons in the substantia nigra – succumb to degeneration. As previously suggested ^68^ for primary neurons seeded with PFFs, toxicity might be conferred to the aggregations in presence of additional insult factors or after more advanced maturation (into LBs).

### Correlation studies of behavior and pS129 immunoreactivity

The finding of small effects of α-Syn pathology on amygdala physiology prompted us to further investigate potential correlations of α-Syn pathology in other brain regions with behavior and physiological parameters. We determined several such parameters by discriminant analysis, differentiating the PFF(C) condition better than the PFF condition from controls. This supports again the notion, that α-Syn pathology by itself does not strongly impact on behavior. To investigate potential correlations of α-Syn pathology in distinct brain regions with behavioral outcomes more closely, we created correlation matrices for behavioral and physiological outcomes with α-Syn pathology in specific brain regions. pS129 immunoreactivity in the entorhinal cortex for PFF-injected –including PFF-injected CORT-treated–mice was negatively correlated with feeding, drinking, hedonic behavior and movement in the EPM, whereas positive correlations were observed for body fat content and depressive-like behavior in the FST, which is interesting in the light of anti-depressive effects demonstrated by activation of the entorhinal cortex^69^. Correlation patterns for FST and AC parameters were intriguingly inverted in the visual, as compared to the entorhinal cortex. Substantia nigra α-Syn pathology was correlated with motor behaviors: negatively with distance travelled in the AC (similar effects of somatosensory and cingulate cortices, but – surprisingly – positively with performance on the Rotarod. α-Syn pathology in the amygdala coincided with reduced explorative behavior in the EPM and with reduced drinking. These results were essentially confirmed by PLS regression analysis, which strikingly separated PFF and PFF(C) groups in all analyzed brain regions very clearly; in this model regarding α-Syn pathology in the substantia nigra, we identified additionally increased associated fear behaviors (freezing in the cue and context test), (non-aggressive) sniffing behavior in the resident intruder test and anxiety like behavior in the OF (latency to enter the center) as more positively correlated with CORT treatment. Interestingly, pS129 immunoreactivity in the prelimbic cortex, which was among the highest among all brain regions; was differentially associated with aggressive and fear behaviors (more negatively associated in PFF(C) mice), as well as with body fat content and weight gain (more positively associated in PFF(C) mice). Similar to the effect of the reversal of hypo-anxiety-like behavior in the EPM and OF by CORT treatment, these correlative results demonstrate divergent physiological and behavioral alterations in heightened CORT conditions in PFF-injected mice, potentially mediated by CORT-suppressed beneficial adaptations (shift to complex-1 linked respiration, hypo-anxiety), leading to higher vulnerability of PFF(C) mice to α-Syn pathology and neurodegeneration. These adaptations did not appear to be linked to differential neuroinflammation, as increased astrogliosis was only observed at the site of injection of PFF and was not modulated by CORT (suppl. Fig. 17).

## Conclusions

We report aggravated α-Syn pathology spreading and neurodegeneration in mice injected with α-Syn PFFs, in a condition of heightened CORT. This suggests heightened glucocorticoid levels risk factor for the development of the neuropathological hallmarks of PD and potential target for treatment. Taken together, our findings suggest that chronic CORT-treatment reduces the ability of the mouse brain to adapt to the additional proteostatic stress of intrastriatal injection of α-Syn PFFs, resulting in lower thresholds for α-Syn pathology handling and nigral neurodegeneration. Further elucidation of (molecular) vulnerability factors of specific brain regions to α-Syn pathology, and why at some point resilience fails and neurodegeneration (such as in the substantia nigra) occurs, will be of importance to understand the complex effects of α-Syn pathology. Based on our results, we suggest that α-Syn pathology in absence of additional clinical (such as depression, chronic stress and potentially anxiety, sleep disturbances, etc) and molecular (reduced mitochondrial function, reduced anti-oxidative capacities, etc.) risk factors is not immediately noxious, maybe even triggering transient protective adaptations. This is in line with reported inconsistencies of LB-pathology and clinical symptoms in human patients and moderate reflection of general PD-symptomatology in α-Syn-based rodent models ^70^.

## Acknowledgments

We are grateful to Ioannis Zalachoras, Laia Morató Fornaguera, Anne Michel, Georges Mairet-Coello, Rachel Angers, Patrick Downey and Martin Citron for valuable intellectual inputs. We thank the following core facilities of the EPFL for excellent technical assistance: Phenotyping Unit (UDP), Histology Core Facility (HCF) and Bioimaging and Optics Platform (BIOP). This work was funded by UCB S.A. and EPFL.

## Author contributions

*JB*: Conceptualization, Data curation, Formal analysis, Investigation, Writing-original draft, *JCC:* Conceptualization, Data curation, Formal analysis, Investigation, *JR:* Formal analysis, *SKT, AC, IGS:* Data curation, Formal analysis, CS: Conceptualization, Formal analysis, Writing – original draft, *HAL:* Conceptualization, Formal analysis, Funding acquisition, Writing – original draft

All authors reviewed, edited and approved of the manuscript.

## Materials and methods

### Preparation and Characterization of PFFs

α-Syn PFFs were generated from recombinant mouse (m) α-Syn protein. The lyophilized protein was dissolved in PBS at a concentration of 325 μM and set to pH7.2. The solution was centrifuged for 5 min through a 0.2 μM filter at 5000 rpm and purity was confirmed by mass spectrometry and HPLC. Supernatant was incubated under constant agitation of 900 rpm on an orbital shaker at 37°C for 4 days. The generated mα-Syn fibrils were sonicated briefly (40% amplitude, one pulse for 5 s) and then were aliquoted and stored at −80 °C.

Thioflavin T (ThT) binding was performed to assess amyloid formation with excitation at 450 nm, emission at 485 nm (Bucher Analyst AD plate reader). Samples were treated with ThT (10 μM, in 50 mM glycine, pH8.5) in black 384-well plates (Nunc).

Remaining soluble protein was assessed by sedimentation assay (supernatant after centrifugation at 100000g for 30 min) and filtration assay (14000 g for 15 min through a 100 kD filter) and analysed by SDS-PAGE (15% polyacrylamide gel) and Coomassie (Life Technologies) staining.

Samples were applied on glow-discharged Formvar/carbon-coated 200-mesh copper grids for analysis by transmission electron microscopy.

### Animals and surgical procedure

C57BL/6JRj male mice were ordered at an age of 8 weeks (Elevage Janvier) and allowed to acclimate to the animal house for at least 2 weeks. They were kept at 23 °C (40 % humidity) in a 12h/12h light/dark cycle (7am-7pm and 7pm-7am, respectively) and free access to standard laboratory rodent chow and water, 3 animals per cage. For the resident intruder test younger BalbC mice (10-12 weeks at test; 3 per cage) and female C57BL/6JRj (3 months old at test) were purchased 2 week before the test.

All animal experimentation procedures were approved by the Cantonal Veterinary Authorities (Vaud, Switzerland) and performed in compliance with the European Communities Council Directive of 24 November 1986 (86/609EEC). Every effort was taken to minimize the number of animals used.

Surgical procedures were performed at an age of 3-5. Wild-type α-Syn PFFs were stereotaxically injected unilaterally into the right dorsal striatum (coordinates: AP +0.4, ML +2, DV-2,6). Fully anesthetized animals (100 mg/kg ketamine and 10 mg/kg xylazine, i.p.) were mounted on stereotactic frames (Kopf Instruments), lubricant eye ointment (Viscotears) was applied and 5 μg PFFs in 2 μL PBS were injected using a 10 μL Hamilton syringe attached to a 34-gauge canula at a flow rate of 0.1 ul/min. Skin incisions were sutured with dissolvable stitches (Vicryl 6.0). All animals were monitored until fully awake and treated with paracetamol (200-300 mg/kg; 2 mg/ml) in the water bottle for 3 days after surgery.

After behavioural experiments (N=9 per group), animals were killed by an overdose of thiopental (150 mg/kg) and after removing blood with heparinized saline (0.9%) brains were fixed by transcardial perfusion with 4 % paraformaldehyd for immunohistochemistry and histological studies. Mice for mitochondrial respiration studies (N=8-9 per group) were killed by neck dislocation and used immediately to measure oxidative phosphorylation parameters.

### Assessment of body composition

Body weight of animals was continuously assessed throughout the experiments. Fat- and lean-mass was measured by Echo MRI at start of corticosterone/vehicle treatment (considered for randomization to experimental groups) and 3 weeks after surgery.

### Continuous exogenous corticosterone treatment

Corticosterone (CORT, Sigma) was dissolved in 0.45 % hydroxypropyl-b-cyclodextrin (Sigma). Either corticosterone (35 mg/l) in hydroxypropyl-b-cyclodextrin or hydroxypropyl-b-cyclodextrin alone (vehicle) was administered to animals in drinking water starting 4 weeks before surgery and then continuously until sacrifice of the animals as described elsewhere (Bacq et a I, 2012).

### Behavioral tests

Starting 4 weeks before surgery, animals were handled weekly (removed from the cage gently, occasionally weighed) until start of behavioural testing and weighed weekly until the end of behavioural tests. All behavioural tests were performed in the morning (8am-lpm), unless stated otherwise. Camcorders (Sony) were used to record behaviour, where applicable.

#### Elevated plus maze

Mice were habituated to the experimental room for at least 45min. They were then placed in the central area of an elevated plus maze (65 cm above the floor, with 2 open and 2 enclosed arms) and allowed to explore the maze for 5 minutes. Maze was cleaned with 5% ethanol between runs. Exploratory behaviour and the time spent in each arm or the center was recorded. Lux in distal parts of open arms was 12.1-12.2 and 8.7 at the mid junction. Ethovision-software (Noldus) was used to score behavior.

#### Open field and novel object test

Light intensity was adjusted to 7 lux in the center of squared boxes. Mice were habituated to the experimental room for more than 30 min before the test and were then placed in the open-field arena. After 10 min a novel object (transparent drinking bottle) was placed in the middle of the arena and mice were again allowed to explore freely for 5 min. Distance travelled and the time spent in the different areas defined in the arena (wall, intermediate and center) were recorded. Ethovision-software (Noldus) was used to score behavior.

#### Marble burying test

Experimental cages were filled approximately 4-5 cm high with bedding. Mice were habituated to the testing room at least 30 minutes before the test. For 15 min, mice were allowed to explore the experimental cage (in absence of marbles). Mice were removed from the cages, 12 glass marbles were placed per cage in a regular pattern on the surface of bedding material, evenly spaced around 4 cm apart, after which the mouse was put back into the cage for 45 minutes. The number of visible marbles was assessed throughout the time of the experiment.

#### Fear conditioning

Mice were habituated to the conditioning chamber for at least for 30 min before each experiment.

(1) Fear acquisition was performed in transparent conditioning chambers equipped with metal grids on a floor plate and cleaned before each trial with 5% ethanol. 3 min after a mouse was placed into the chamber, it was exposed to a 30 s (800 Hz, 80 dB) long auditory cue (Acoustic Stimuli LE 114, Panlab, s.I.), followed by a 2 s, 0.5 mA electrical foot shock (Shocker LE 100-26, Letica) delivered via the metal grid. 1 and 2.5 min later auditory cue and foot shock were repeated.

(2) Context test was performed 1 day later in the same context as in fear acquisition. Mice were placed for 7.5 min in the conditioning chamber without auditory cues or foot shocks being delivered. Chambers were cleaned before each trial with 5% ethanol.

(3) Extinction training was performed on the same day in the afternoon in a new context: room light was dimmed, the floor grid was replaced by a white floor plate, 5% vanilla-solution was used for cleaning. A circular cage with jungle-like painting was used instead of the transparent chambers. 3 min after a mouse was placed in the conditioning chamber, it was exposed to 12 consecutive 30 s long auditory cues (same as in fear acquisition), interspersed by silent phases of 1 min each.

(4) In the next morning, mice were placed in the same context as for extinction training, 5% vanilla-solution was used for cleaning. 3 min after a mouse was placed into the chamber, it was exposed to 3 consecutive 30 s long auditory cues (same as in fear acquisition), interspersed by silent phases of 1 min each.

Softwares *Freezing vl.3.05* and *The Observer XT 11.5* were used to program the different protocols and to analyze fear behavior, respectively. Freezing was defined as the absence of all movement, except for breathing. All sequences with tone exposure were scored for acquisition, extinction and tone test. Context test was scored entirely.

#### Forced swim test

5L glass beakers were filled with 3-3.5 L tap water (25°C). Mice were exposed to the water for 15 min on day 1 and another 5 min on the following day. *Ethovision* (Noldus) software was used to quantify immobile floating versus active swimming behavior.

#### Resident intruder test

Tested male mice (residents) were housed in a standard cage with a female for 4 days prior to the test to facilitate the development of territoriality. The bedding of the cage was not changed during that initial period. On the day of the test, the female was removed from the cage. 30 min later, an unfamiliar younger, male BalbC (intruder) with 5-15% less weight was introduced into the home cage. The two mice were allowed to interact physically for 10 min.

Resident intruder tests were performed exclusively within the dark period of the animal house (between 7pm and 1am).

#### Rotarod

Animals were habituated to the experimental room for at least 30 min. Mice were placed on the lanes of a Rotarod apparatus (BIOSEB) rod with an empty lane between each mouse. During training and testing, the same mouse was always put on the same lane. Acceleration speed was set to 4-40 rpm. 3 trials were realized for each mouse, with an inter-trial break of 15 min. Maximum trial duration was 300 s. The apparatus was cleaned between each trial with 5% mucosol.

On day 1 animals were trained to stay on the rotarod rod. Training was stopped in case of three consecutive times of the following events: passive rotation, jumping off, or falling off the rod.

Animals were tested 1 day later, 3 trials each. Each test was ended, when the mouse passively rotated, jumped or fell off the rod or if 300 s elapsed. For analysis, only day 2 was used. The average of the best 2 trials were used for analysis.

#### Activity cage and saccharin preference

Mice were single caged in an activity cage (TSE-systems) for 72 h. By means of a two-bottle preference test intake of saccharin solution (0.05% saccharin sodium salt, Sigma) was compared with intake of tap water. Additionally, overall drinking and food consumption of the mice was recorded and general motor activity was assessed. All parameters were recorded in 30 min intervals.

### Respirometry

Mice for respirometric experiments were sacrificed by neck dislocation and amygdala and striatum were dissected on ice using a mouse brain matrix (Agnthos). Wet tissue was weighed and collected in ice-cold BIOPS (2.8 mM Ca_2_K_2_EGTA, 7.2 mM K_2_EGTA, 5.8 mM ATP, 6.6 mM MgCl_2_, 20 mM taurine, 15 mM sodium phosphocreatine, 20 mM imidazole, 0.5 mM dithiothreitol and 50 mM MES, pH = 7.1), homogenized in ice-cold MiR05 (0.5 mM EGTA, 3mM MgCl_2_, 60 mM potassium lactobionate, 20 mM taurine, 10 mM KH_2_PO_4_, 20 mM HEPES, 110 mM sucrose and 0.1% (w/v) BSA, pH=7.1) using a pestle for eppendorf tubes in a concentration of 1 mg wet-weight per 10 μL MiR05. Respiration was measured in parallel to mitochondrial ROS production (O_2_^−^ and H_2_O_2_) at 37 °C in the Oroboros 02k equipped with the 02K Fluo-LED2 Module (Oroboros Instruments, Austria). For mitochondrial ROS-measurement LEDs for green excitation were applied and a concentration of 1 mg wet tissue per ml MiR05 was added to final concentrations of 10 μM amplex red, 1 U/ml horse radish peroxidase and 5 U/ml superoxide dismutase in 2 ml MiR05 per 02K chamber. Calibration was performed by titrations of 5 μL of 40 μM H_2_O_2_.

A substrate-uncoupler-inhibitor-titration (SUIT) protocol was applied to measure oxygen flux at different repirational states as described previously (Hollis et al, 2015; Burtscher et al, 2015). Briefly, NADH-pathway (N) respiration in the LEAK and oxidative phosphorylation (OXPHOS) state was analysed in presence of malate (2mM), pyruvate (lOmM) and glutamate (2OmM) before and after the addition of ADP (5 mM), respectively (N*_L_*, N*_P_*). Addition of succinate (10 mM) allowed assessment of NADH- and Succinate-linked respiration in OXPHOS (NS*_P_*) and in the uncoupled state (NS*_E_*) after incremental (Δ0.5 μM) addition of carbonyl cyanide m-chlorophenyl hydrazine (CCCP). Inhibition of Complex I by rotentone (0.5 μM) yielded succinate-linked respiration in the uncoupled state (S*_E_*). Tissue-mass specific oxygen fluxes were corrected for residual oxygen consumption, *Rox*, measured after additional inhibition of the mitochondrial electron transfer system, ETS, Complex III with antimycin A. For further normalization, fluxes of all respiratory states were divided by ET-capacity to obtain flux control ratios, *FCR*. Terminology was applied according to http://www.mitoeagle.org/index.php/MitoEAGLE_preprint_2018-02-08.

### Brain tissue preparation for gel electrophoresis and western blots

Brain homogenates not used for respirometry were treated with protease and phosphatase inhibitors, snap-frozen and stored at −80°C. Protein concentrations were determined by BCA assay (Thermo Scientific), samples were diluted in 4x Laemmli buffer, boiled for 10 min and separated on a 15% SDS-PAGE gel and transferred onto nitrocellulose membrane (Fisher Scientific, Lucens, Switzerland) with a semi-dry system (Bio-Rad, Crissier, Switzerland). Membranes were probed overnight at 4?°C with the primary antibody of interest after 30?min of blocking in Odyssey blocking buffer (Li-Cor Biosciences, Bad Homburg, Germany) diluted 113:133 in PBS. After four washes with PBS and 0.01% (v/v) Tween-20 (PBS-T), membranes were incubated for 1?h with secondary antibodies (goat Alexa Fluor^680^ IgG) protected from light at RT. Immunoblots were finally washed 4 times with PBS-T and scanned using a Li-COR scanner (Li-Cor Biosciences) at a wavelength of 700?nm. Image J was used for densitometry.

### Immunohistochemistry and imaging

Immunohistochemistry and Mayer’s hematoxylin stainings were performed on sections of brains fixed in 4% paraformaldehyde, embedded in paraffin and cut coronally to 4 μm. Sections were de-waxed and epitope retrieval was performed for 20 min at 95°C in trisodium citrate buffer (10mM, pH 6.0) in a retriever (Labvision). Sections were then blocked for 60 min in 3% bovine serum albumin in PBS containing 0.1% Triton X-100 at RT. Primary antibodies were applied over night at 4°C and secondary antibodies for immunofluorescence for 60 min at RT, before mounting the slides using fluoromount. In case of 3,3’-diaminobenzidine (DAB) – revelation, sections were exposed to 3% H_2_O_2_ in PBS for 30 min before blocking and ImmPRESS reagent anti-mouse IgG (Vector MP-7402) or anti-goat IgG (Vector MP-7405) was applied for 40 min at RT instead of fluorescent secondary antibodies, followed by incubation for 10 min in DAB dissolved in 50 mM Tris buffer and 0.06% H_2_O_2_. Sections were counterstained with Mayer’s hematoxylin and mounted with fluoromount. In case of Proteinase K (PK) treatment, sections were incubated for 8 min at room temperature in 1 μg/mL of PK in 5OmM TrisHCI buffer (pH 7.4).

Tiled images for TH were produced with a Leica DM5500 microscope, an Olympus AX70 microscope was used to create images from DAB-stained sections and Zeiss LSM700 confocal microscopes or Leica DM5500 were used to image immunofluorescent sections. Image J was used to assess densities of immunoreactivity (coverage of area with immunoreactivity in % of total area).

**Table 1.** antibodies employed

| type | species | specification | Concentration | application |
| --- | --- | --- | --- | --- |
| aSyn pS129 | mouse | Wako 014-20281 | 1:10000<br>(DAB), 1:1000<br>(IF) | IHC |
| Tyrosine hydroxylase | rabbit | Millipore AB152 | 1:1000 | IHC |
| MJF-R13 | rabbit | Abcam 168381 | 1:750 | IHC |
| MAP2 | chicken | Abcam ab5392 | 1:2000 | IHC |
| GFAP | goat | Santa Cruz sc-6170 | 1:500 | IHC |
| Ubiquitin 1 | mouse | Millipore MAB1510 | 1:500 | IHC |
| p62 | mouse | Abcam ab56416 | 1:1000 | IHC |
| aSyn 1-20 | rabbit | Eurogentec | 1:1000 | IHC |
| antiRabbit 647 | donkey | Invitrogen | 1:800 | IHC |
| antiMouse 488 | goat | Invitrogen | 1:800 | IHC |
| antiMouse 568 | goat | Invitrogen | 1:800 | IHC |
| antiChicken 488 | donkey | Jackson lab | 1:500 | IHC |
| ImmPRESS antiMouse | horse | Vector MP-7402 | 1 drop /<br>section | IHC |
| ImmPRESS antiGoat | horse | Vector MP-7405 | 1 drop /<br>section | IHC |
| Synuclein (SYN1) | mouse | BD Transduction,<br>BD610787 | 1:1000 | Western<br>blot |
| Beta-actin | mouse | Abcam Ab6276-100 | 1:5000 | Western<br>blot |
| anti-mouse 680 IRDye<br>680RD | goat | Li-COR 926-68070 | 1:10000 | Western<br>blot |

### Statistical analyses

Data are presented as means ± SD, except for behavioural tests, in which means ± SEM are presented. Heat maps are based on mean values (of pS129 immunodensities for brain regions). Statistical tests applied for the different experiments are given in figure legends. P values <0.05 were considered as significant. Pearson coefficients were calculated for correlation studies. Microsoft Office Excel and Graphpad Prism were used to present statistical results, except for PLS-analyses (see below).

#### PLS Discriminant analysis

Partial least squares discriminant analysis (PLS-DA) was used to determine which of the behavioral and physiological variables assessed in this study best discriminate among PFF injected groups (CORT and vehicle) and PBS-injected controls (vehicle). For each variable, mean value imputation was used, missing scaling was performed by mean centering and dividing by the square root of the standard deviation (z-scoring).

The cross validation (CV) procedure performed was the 10-fold CV, with prediction accuracy as the measured performance metric. PLS-DA model validation relied on permutation tests using 2000 permutations where, for each permutation, a PLS-DA model is built for the data with permuted group labels, and its prediction accuracy is calculated. The null-hypothesis of a nonsignificant discriminant model is rejected if the prediction accuracy with the original groups is not a part of the distribution based on the permuted group assignment (above the 95^th^ percentile).

Analysis was performed in MetaboAnalyst 4.0 (http://www.metaboanalyst.ca)

#### PLS Regression analysis

Partial least squares (PLS) regression was used to analyze levels of α-Syn pS129 immunoreactivity for PFF-injected groups in pre-selected brain regions behavior variables. For each variable, mean value imputation was used, missing scaling was performed by mean centering and dividing by the square root of the standard deviation (z-scoring). For each brain region, the resulting PLS components with a percentage of variance explained larger than 15% were selected and used as the dependent variable in a multivariate analysis of variance (MANOVA) using the PFF(C) group as the independent variable and all the pre-selected components as dependent variables. Follow-up ANOVAs based on a significant MANOVA outcome were corrected for multiple comparisons (Holm–Bonferroni method). PLS regression was implemented with the Matlab (version 2018a) function *pisregress*.

## References

1 Spillantini, M. G. et al. Alpha-synuclein in Lewy bodies. Nature 388, 839–840, doi:10.1038/42166 (1997).

2 Lashuel, H. A., Overk, C. R., Oueslati, A. & Masliah, E. The many faces of alpha-synuclein: from structure and toxicity to therapeutic target. Nature reviews. Neuroscience 14, 38–48, doi:10.1038/nrn3406 (2013).

3 Burre, J. et al. Alpha-synuclein promotes SNARE-complex assembly in vivo and in vitro. Science (New York, N.Y.) 329, 1663–1667, doi:10.1126/science.1195227 (2010).

4 Abeliovich, A. et al. Mice lacking alpha-synuclein display functional deficits in the nigrostriatal dopamine system. Neuron 25, 239–252 (2000).

5 Logan, T., Bendor, J., Toupin, C., Thorn, K. & Edwards, R. H. alpha-Synuclein promotes dilation of the exocytotic fusion pore. Nature neuroscience 20, 681–689, doi:10.1038/nn.4529 (2017).

6 Ludtmann, M. H. et al. Monomeric Alpha-Synuclein Exerts a Physiological Role on Brain ATP Synthase. The Journal of neuroscience : the official journal of the Society for Neuroscience 36, 10510–10521, doi:10.1523/jneurosci.1659-16.2016 (2016).

7 Polymenidou, M. & Cleveland, D. W. Prion-like spread of protein aggregates in neurodegeneration. The Journal of experimental medicine 209, 889–893, doi:10.1084/jem.20120741 (2012).

8 Polymeropoulos, M. H. et al. Mutation in the alpha-synuclein gene identified in families with Parkinson’s disease. Science (New York, N.Y.) 276, 2045–2047 (1997).

9 Kruger, R. et al. Ala30Pro mutation in the gene encoding alpha-synuclein in Parkinson’s disease. Nature genetics 18, 106–108, doi:10.1038/ng0298-106 (1998).

10 Zarranz, J. J. et al. The new mutation, E46K, of alpha-synuclein causes Parkinson and Lewy body dementia. Annals of neurology 55, 164–173, doi:10.1002/ana.10795 (2004).

11 Singleton, A. B. et al. alpha-Synuclein locus triplication causes Parkinson’s disease. Science (New York, N.Y.) 302, 841, doi:10.1126/science.1090278 (2003).

12 Ibanez, P. et al. Causal relation between alpha-synuclein gene duplication and familial Parkinson’s disease. Lancet (London, England) 364, 1169–1171, doi:10.1016/s0140-6736(04)17104-3 (2004).

13 Dawson, T. M., Golde, T. E. & Lagier-Tourenne, C. Animal models of neurodegenerative diseases. Nature neuroscience 21, 1370–1379, doi:10.1038/s41593-018-0236-8 (2018).

14 Kalia, L. V. & Lang, A. E. Parkinson’s disease. Lancet (London, England) 386, 896–912, doi:10.1016/s0140-6736(14)61393-3 (2015).

15 Shiba, M. et al. Anxiety disorders and depressive disorders preceding Parkinson’s disease: a case-control study. Movement disorders : official journal of the Movement Disorder Society 15, 669–677 (2000).

16 Sagna, A., Gallo, J. J. & Pontone, G. M. Systematic review of factors associated with depression and anxiety disorders among older adults with Parkinson’s disease. Parkinsonism & related disorders 20, 708–715, doi:10.1016/j.parkreldis.2014.03.020 (2014).

17 Thobois, S., Prange, S., Sgambato-Faure, V., Tremblay, L. & Broussolle, E. Imaging the Etiology of Apathy, Anxiety, and Depression in Parkinson’s Disease: Implication for Treatment. Current neurology and neuroscience reports 17, 76, doi:10.1007/s11910-017-0788-0 (2017).

18 Kavushansky, A. & Richter-Levin, G. Effects of stress and corticosterone on activity and plasticity in the amygdala. Journal of neuroscience research 84, 1580–1587, doi:10.1002/jnr.21058 (2006).

19 Janak, P. H. & Tye, K. M. From circuits to behaviour in the amygdala. Nature 517, 284–292, doi:10.1038/nature14188 (2015).

20 Popescu, A., Lippa, C. F., Lee, V. M. & Trojanowski, J. Q. Lewy bodies in the amygdala: increase of alpha-synuclein aggregates in neurodegenerative diseases with tau-based inclusions. Archives of neurology 61, 1915–1919, doi:10.1001/archneur.61.12.1915 (2004).

21 Nelson, P. T. et al. The Amygdala as a Locus of Pathologic Misfolding in Neurodegenerative Diseases. Journal of neuropathology and experimental neurology 77, 2–20, doi:10.1093/jnen/nlx099 (2018).

22 Kordower, J. H., Chu, Y., Hauser, R. A., Freeman, T. B. & Olanow, C. W. Lewy body-like pathology in long-term embryonic nigral transplants in Parkinson’s disease. Nature medicine 14, 504–506, doi:10.1038/nm1747 (2008).

23 Li, J. Y. et al. Lewy bodies in grafted neurons in subjects with Parkinson’s disease suggest host-to-graft disease propagation. Nature medicine 14, 501–503, doi:10.1038/nm1746 (2008).

24 Desplats, P. et al. Inclusion formation and neuronal cell death through neuron-to-neuron transmission of alpha-synuclein. Proceedings of the National Academy of Sciences of the United States of America 106, 13010–13015, doi:10.1073/pnas.0903691106 (2009).

25 Volpicelli-Daley, L. A. et al. Exogenous alpha-synuclein fibrils induce Lewy body pathology leading to synaptic dysfunction and neuron death. Neuron 72, 57–71, doi:10.1016/j.neuron.2011.08.033 (2011).

26 Mougenot, A. L. et al. Prion-like acceleration of a synucleinopathy in a transgenic mouse model. Neurobiology of aging 33, 2225–2228, doi:10.1016/j.neurobiolaging.2011.06.022 (2012).

27 Luk, K. C. et al. Pathological alpha-synuclein transmission initiates Parkinson-like neurodegeneration in nontransgenic mice. Science (New York, N.Y.) 338, 949–953, doi:10.1126/science.1227157 (2012).

28 Masuda-Suzukake, M. et al. Prion-like spreading of pathological alpha-synuclein in brain. Brain : a journal of neurology 136, 1128–1138, doi:10.1093/brain/awt037 (2013).

29 Rey, N. L., George, S. & Brundin, P. Review: Spreading the word: precise animal models and validated methods are vital when evaluating prion-like behaviour of alpha-synuclein. Neuropathology and applied neurobiology 42, 51–76, doi:10.1111/nan.12299 (2016).

30 Recasens, A. et al. Lewy body extracts from Parkinson disease brains trigger alpha-synuclein pathology and neurodegeneration in mice and monkeys. Annals of neurology 75, 351–362, doi:10.1002/ana.24066 (2014).

31 Prusiner, S. B. et al. Evidence for alpha-synuclein prions causing multiple system atrophy in humans with parkinsonism. Proceedings of the National Academy of Sciences of the United States of America 112, E5308–5317, doi:10.1073/pnas.1514475112 (2015).

32 Peng, C. et al. Cellular milieu imparts distinct pathological alpha-synuclein strains in alpha-synucleinopathies. Nature, doi:10.1038/s41586-018-0104-4 (2018).

33 Rey, N. L. et al. Widespread transneuronal propagation of alpha-synucleinopathy triggered in olfactory bulb mimics prodromal Parkinson’s disease. The Journal of experimental medicine 213, 1759–1778, doi:10.1084/jem.20160368 (2016).

34 Sandi, C. & Richter-Levin, G. From high anxiety trait to depression: a neurocognitive hypothesis. Trends in neurosciences 32, 312–320, doi:10.1016/j.tins.2009.02.004 (2009).

35 de Kloet, E. R. et al. Stress and Depression: a Crucial Role of the Mineralocorticoid Receptor. Journal of neuroendocrinology 28, doi:10.1111/jne.12379 (2016).

36 Hemmerle, A. M., Dickerson, J. W., Herman, J. P. & Seroogy, K. B. Stress exacerbates experimental Parkinson’s disease. Molecular psychiatry 19, 638–640, doi:10.1038/mp.2013.108 (2014).

37 Wu, Q., Yang, X., Zhang, Y., Zhang, L. & Feng, L. Chronic mild stress accelerates the progression of Parkinson’s disease in A53T alpha-synuclein transgenic mice. Experimental neurology 285, 61–71, doi:10.1016/j.expneurol.2016.09.004 (2016).

38 Gourley, S. L., Kiraly, D. D., Howell, J. L., Olausson, P. & Taylor, J. R. Acute hippocampal brain-derived neurotrophic factor restores motivational and forced swim performance after corticosterone. Biological psychiatry 64, 884–890, doi:10.1016/j.biopsych.2008.06.016 (2008).

39 David, D. J. et al. Neurogenesis-dependent and -independent effects of fluoxetine in an animal model of anxiety/depression. Neuron 62, 479–493, doi:10.1016/j.neuron.2009.04.017 (2009).

40 Bacq, A. et al. Organic cation transporter 2 controls brain norepinephrine and serotonin clearance and antidepressant response. Molecular psychiatry 17, 926–939, doi:10.1038/mp.2011.87 (2012).

41 Spillantini, M. G., Crowther, R. A., Jakes, R., Hasegawa, M. & Goedert, M. alpha-Synuclein in filamentous inclusions of Lewy bodies from Parkinson’s disease and dementia with lewy bodies. Proceedings of the National Academy of Sciences of the United States of America 95, 6469–6473 (1998).

42 Carulla, N. et al. Molecular recycling within amyloid fibrils. Nature 436, 554–558, doi:10.1038/nature03986 (2005).

43 Mahul-Mellier, A. L. et al. Fibril growth and seeding capacity play key roles in alpha-synuclein-mediated apoptotic cell death. Cell death and differentiation 22, 2107–2122, doi:10.1038/cdd.2015.79 (2015).

44 Rebuffe-Scrive, M., Walsh, U. A., McEwen, B. & Rodin, J. Effect of chronic stress and exogenous glucocorticoids on regional fat distribution and metabolism. Physiology & behavior 52, 583–590 (1992).

45 Rogan, M. T., Staubli, U. V. & LeDoux, J. E. Fear conditioning induces associative long-term potentiation in the amygdala. Nature 390, 604–607, doi:10.1038/37601 (1997).

46 Coccaro, E. F., McCloskey, M. S., Fitzgerald, D. A. & Phan, K. L. Amygdala and orbitofrontal reactivity to social threat in individuals with impulsive aggression. Biological psychiatry 62, 168–178 (2007).

47 Di Maio, R. et al. alpha-Synuclein binds to TOM20 and inhibits mitochondrial protein import in Parkinson’s disease. Science translational medicine 8, 342ra378, doi:10.1126/scitranslmed.aaf3634 (2016).

48 Grassi, D. et al. Identification of a highly neurotoxic alpha-synuclein species inducing mitochondrial damage and mitophagy in Parkinson’s disease. Proceedings of the National Academy of Sciences of the United States of America 115, E2634–E2643, doi:10.1073/pnas.1713849115 (2018).

49 Tapias, V. et al. Synthetic alpha-synuclein fibrils cause mitochondrial impairment and selective dopamine neurodegeneration in part via iNOS-mediated nitric oxide production. Cellular and molecular life sciences : CMLS 74, 2851–2874, doi:10.1007/s00018-017-2541-x (2017).

50 Ludtmann, M. H. R. et al. alpha-synuclein oligomers interact with ATP synthase and open the permeability transition pore in Parkinson’s disease. Nature communications 9, 2293, doi:10.1038/s41467-018-04422-2 (2018).

51 Castrioto, A., Thobois, S., Carnicella, S., Maillet, A. & Krack, P. Emotional manifestations of PD: Neurobiological basis. Movement disorders : official journal of the Movement Disorder Society 31, 1103–1113, doi:10.1002/mds.26587 (2016).

52 Campos, F. L. et al. Rodent models of Parkinson’s disease: beyond the motor symptomatology. Frontiers in behavioral neuroscience 7, 175, doi:10.3389/fnbeh.2013.00175 (2013).

53 Caudal, D., Alvarsson, A., Bjorklund, A. & Svenningsson, P. Depressive-like phenotype induced by AAV-mediated overexpression of human alpha-synuclein in midbrain dopaminergic neurons. Experimental neurology 273, 243–252, doi:10.1016/j.expneurol.2015.09.002 (2015).

54 Janakiraman, U. et al. Influences of Chronic Mild Stress Exposure on Motor, Non-Motor Impairments and Neurochemical Variables in Specific Brain Areas of MPTP/Probenecid Induced Neurotoxicity in Mice. PloS one 11, e0146671, doi:10.1371/journal.pone.0146671 (2016).

55 Mattila, P. M., Rinne, J. O., Helenius, H., Dickson, D. W. & Roytta, M. Alpha-synucleinimmunoreactive cortical Lewy bodies are associated with cognitive impairment in Parkinson’s disease. Acta neuropathologica 100, 285–290 (2000).

56 Jellinger, K. A. Alpha-synuclein pathology in Parkinson’s and Alzheimer’s disease brain: incidence and topographic distribution--a pilot study. Acta neuropathologica 106, 191–201, doi:10.1007/s00401-003-0725-y (2003).

57 Gómez-Tortosa, E., Newell, K., Irizarry, M. C., Sanders, J. L. & Hyman, B. T. α-Synuclein immunoreactivity in dementia with Lewy bodies: morphological staging and comparison with ubiquitin immunostaining. Acta neuropathologica 99, 352–357 (2000).

58 Kovari, E. et al. Lewy body densities in the entorhinal and anterior cingulate cortex predict cognitive deficits in Parkinson’s disease. Acta neuropathologica 106, 83–88, doi:10.1007/s00401-003-0705-2 (2003).

59 George, S. et al. Alpha-synuclein transgenic mice exhibit reduced anxiety-like behaviour. Experimental neurology 210, 788–792, doi:10.1016/j.expneurol.2007.12.017 (2008).

60 Graham, D. R. & Sidhu, A. Mice expressing the A53T mutant form of human alpha-synuclein exhibit hyperactivity and reduced anxiety-like behavior. Journal of neuroscience research 88, 1777–1783, doi:10.1002/jnr.22331 (2010).

61 Ardouin, C. et al. [Assessment of hyper- and hypodopaminergic behaviors in Parkinson’s disease]. Revue neurologique 165, 845–856, doi:10.1016/j.neurol.2009.06.003 (2009).

62 Nemani, V. M. et al. Increased expression of alpha-synuclein reduces neurotransmitter release by inhibiting synaptic vesicle reclustering after endocytosis. Neuron 65, 66–79, doi:10.1016/j.neuron.2009.12.023 (2010).

63 Froula, J. M. et al. alpha-Synuclein fibril-induced paradoxical structural and functional defects in hippocampal neurons. Acta neuropathologica communications 6, 35, doi:10.1186/s40478-018-0537-x (2018).

64 Greene, J. G., Porter, R. H., Eller, R. V. & Greenamyre, J. T. Inhibition of succinate dehydrogenase by malonic acid produces an “excitotoxic” lesion in rat striatum. Journal of neurochemistry 61, 1151–1154 (1993).

65 Riepe, M. W. et al. Increased hypoxic tolerance by chemical inhibition of oxidative phosphorylation: “chemical preconditioning”. Journal of cerebral blood flow and metabolism : official journal of the International Society of Cerebral Blood Flow and Metabolism 17, 257–264, doi:10.1097/00004647-199703000-00002 (1997).

66 Lago, T., Davis, A., Grillon, C. & Ernst, M. Striatum on the anxiety map: Small detours into adolescence. Brain research 1654, 177–184, doi:10.1016/j.brainres.2016.06.006 (2017).

67 Nouraei, N. et al. Critical appraisal of pathology transmission in the alpha-synuclein fibril model of Lewy body disorders. Experimental neurology 299, 172–196, doi:10.1016/j.expneurol.2017.10.017 (2018).

68 Fares, M. B. et al. Induction of de novo alpha-synuclein fibrillization in a neuronal model for Parkinson’s disease. Proceedings of the National Academy of Sciences of the United States of America 113, E912–921, doi:10.1073/pnas.1512876113 (2016).

69 Yun, S. et al. Stimulation of entorhinal cortex–dentate gyrus circuitry is antidepressive. Nature medicine 24, 658 (2018).

70 Steiner, J. A., Quansah, E. & Brundin, P. The concept of alpha-synuclein as a prion-like protein: ten years after. Cell and tissue research, doi:10.1007/s00441-018-2814-1 (2018).

